# Neural pattern similarity differentially affects memory performance of younger and older adults: Age differences in neural similarity and memory

**DOI:** 10.1101/528620

**Authors:** Verena R. Sommer, Yana Fandakova, Thomas H. Grandy, Yee Lee Shing, Markus Werkle-Bergner, Myriam C. Sander

## Abstract

Age-related memory decline is associated with changes in neural functioning but little is known about how aging affects the quality of information representation in the brain. Whereas a long-standing hypothesis of the aging literature links cognitive impairments to less distinct neural representations in old age, memory studies have shown that high similarity between activity patterns benefits memory performance for the respective stimuli. Here, we addressed this apparent conflict by investigating between-item representational similarity in 50 younger (19–27 years old) and 63 older (63–75 years old) human adults (male and female) who studied scene-word associations using a mnemonic imagery strategy while electroencephalography was recorded. We compared the similarity of spatiotemporal frequency patterns elicited during encoding of items with different subsequent memory fate. Compared to younger adults, older adults’ memory representations were more similar to each other but items that elicited the most similar activity patterns early in the encoding trial were those that were best remembered by older adults. In contrast, young adults’ memory performance benefited from decreased similarity between earlier and later periods in the encoding trials, which might reflect their better success in forming unique memorable mental images of the joint picture–word pair. Our results advance the understanding of the representational properties that give rise to memory quality as well as how these properties change in the course of aging.

**Significance statement:** Declining memory abilities are one of the most evident limitations for humans when growing older. Despite recent advances of our understanding of how the brain represents and stores information in distributed activation patterns, little is known about how the quality of information representation changes during aging and thus affects memory performance. We investigated how the similarity between neural representations relates to subsequent memory quality in younger and older adults. We present novel evidence that the interaction of pattern similarity and memory performance differs between age groups: Older adults benefited from increased similarity during early encoding whereas young adults benefited from decreased similarity between early and later encoding. These results provide insights into the nature of memory and age-related memory deficits.

## Introduction

A long-standing hypothesis in the cognitive neuroscience of aging holds that neural representations become less specific with advancing age, with detrimental effects on cognitive performance. Reduced neural distinctiveness in older compared to young adults (Li et al., 2001) has been observed as increased similarity and/or reduced discriminability between neural activity patterns during different memory tasks (Carp et al., 2010; St-Laurent et al., 2011), between different stimulus categories (Carp et al., 2011; Park et al., 2004; Park et al., 2010; Park et al., 2012; Payer et al., 2006; Koen et al., 2019), and between different individual stimuli (Goh et al., 2010; St-Laurent et al., 2014). However, these studies did not directly link this age-related reduction in neural specificity to differences in memory performance. A recent functional magnetic resonance imaging (fMRI) study by Koen et al. (2019) assessed neural distinctiveness during a memory encoding task and showed a general (age-invariant) association between neural category differentiation and recognition memory performance. However, they did not identify differences in distinctiveness between items that were later remembered or not remembered. A suitable approach to unravel the specific association between pattern distinctiveness and memory performance would be to examine whether items that are represented less distinctly are also those that are less likely to be remembered. One fMRI study by Zheng et al. (2017) provided first evidence in this direction, showing that decreased memory performance in old age is associated with poorer item-specific representations in the visual cortex, even after controlling for differences in activation levels and variance.

Surprisingly, the hypothesis of the cognitive aging literature suggesting that reduced neural specificity underlies cognitive decline is in stark contrast to the prevalent evidence in general memory research that increased similarity is actually advantageous for performance: In young adult samples, various studies have shown that the representational similarity between different items is positively related to memory performance for these items (Davis et al., 2014; Lu et al., 2015; Wagner et al., 2016), which is in line with cognitive and computational models (Clark and Gronlund, 1996; Gillund and Shiffrin, 1984). Between-item pattern similarity may support memory by capturing regularities across experiences (LaRocque et al., 2013) and by giving rise to a sense of familiarity (Davis et al., 2014).

To date, most studies have used fMRI to estimate neural representations, prioritizing spatial resolution over temporal dynamics of representational patterns. In contrast, time-sensitive magneto-/electroencephalography (M/EEG) measurements are able to identify the precise time windows and processing stages at which representational similarity supports memory performance. Lu et al. (2015) showed that between approximately 420 ms and 580 ms after stimulus onset, global spatiotemporal EEG pattern similarity was higher for later remembered than for forgotten symbols.

Recent scalp (Kerrén et al., 2018; Michelmann et al., 2016) and intracranial EEG studies (Staresina et al., 2016; Zhang et al., 2015) further demonstrated the particular potential of frequency-transformed activity patterns in identifying memory-relevant reactivation of item-specific signatures. For example, Michelmann et al. (2018) showed that desynchronized low-frequency brain oscillations carried stimulus-specific temporal activity patterns during an associative memory task. However, there are no previous reports on the relation of the similarity between these frequency-transformed activation patterns to later memory success for the studied items.

To our knowledge, the apparent conflict between the observed beneficial effect of global similarity in memory studies with young adults, and the potentially detrimental effect of decreasing distinctiveness in the aging literature has not been explicitly addressed. Here, we aim to resolve the question whether distinctiveness or similarity between different neural representations is beneficial for memory performance by a systematic investigation of the relation between representational similarity and memory performance in young and older adults. For this, we examined the similarity of EEG frequency patterns elicited during encoding of scene-word pairs in relation to age and subsequent recall performance.

## Materials and Methods

### Experimental design

The research presented here comprises data from two associated studies that investigated age-related differences in associative memory encoding, consolidation, and retrieval (Fandakova et al., 2018; Muehlroth et al., in press; Sander et al., 2019). Despite subsequent procedural differences, an identical picture–word association task paradigm during which EEG was recorded was at the core of both studies. In this task, participants were asked to memorize scene–word pairs by applying a previously trained mnemonic imagery strategy. Specifically, they were instructed to imagine the scene and word content together in a unique and memorable mental image. Stimuli consisted of color photographs of indoor and outdoor scenes randomly paired with concrete German nouns (4–8 letters). During the initial study phase, scenes and words were presented next to each other on a black background for 4 s. After studying a pair, participants indicated on a four-point scale how well they were able to integrate the presented scene and word. Young and older adults studied 440 and 280 pairs, respectively. During the subsequent cued recall phase, scenes served as cues for participants to verbally recall the associated word. Recall time was not constrained. After each trial, the correct scene–word pair was presented again for 3 s and subjects were instructed to restudy the pair, independent of previous retrieval success. This recall and restudy phase was repeated one more time for the older adults. Finally, both young and older participants underwent a final cued recall round in which no feedback was presented. The number of to-be-studied pairs as well as recall repetitions differed between age groups in order to achieve comparable recall success of approximately half of the studied items. After each phase, we asked participants to indicate on a four-point scale how often they used the instructed imagery strategy or other specific memory strategies to memorize a pair. For a detailed description of the study design and stimulus selection, see Fandakova et al. (2018).

### Subjects

The original sample of study 1 (Fandakova et al., 2018) consisted of 30 healthy young adults and 44 healthy older adults. Due to technical failures, one young adult and three older adults did not complete the study. Study 2 (Muehlroth et al., in press) involved 34 healthy young adults and 41 healthy older adults participated, with 4 younger and 4 older participants not completing the experiment for technical reasons. Due to missing or noisy EEG data, we additionally excluded 9 younger and 15 older adults, resulting in a total of 50 younger adults and 63 older adults across both studies, who are included in the analyses presented here (young adults: *M* (SD)age = 24.3(2.5) years, 19–27 years, 27 female, 23 male; old adults: *M* (SD)age = 70.4(2.6) years, 63–75 years, 33 female, 30 male).

All participants were right-handed native German speakers, reported normal or corrected-to-normal vision, no history of psychiatric or neurological disease, and no use of psychiatric medication. We screened older adults with the Mini-Mental State Examination (MMSE; Folstein et al., 1975) and none had a value below the threshold of 26 points. Both studies were approved by the ethics committee of the Deutsche Gesellschaft für Psychologie and took place at the Max Planck Institute for Human Development in Berlin, Germany. All participants gave written consent to take part in the experiment.

### Behavioral analysis

During the cued recall phases, participants had to verbally recall the word associated with the presented image. We report the proportion of correctly recalled words. False responses occurred rarely and were treated as no responses. Following the rationale of a subsequent memory analysis (Paller and Wagner, 2002), we sorted all trials according to whether the associated word was successfully recalled during the experiment or not. Items that were not remembered after repeated encoding were assumed to have only created a weak memory trace, not sufficient for successful recall (although maybe strong enough for successful recognition, see Fandakova et al., 2018). Importantly, given the repeated recall phases, we were able to further differentiate successfully recalled items and distinguish those that were immediately learned from those that were only acquired later in the experiment. We refer to those items as high memory quality and medium memory quality items, respectively (see Figure 2). Older adults underwent one additional recall and restudy cycle due to close-to-floor performance in the first cycle. To keep the scoring of stimulus pairs as evincing high, medium, or low memory quality comparable across age groups, items that were recalled successfully in the last recall cycle were divided into those that were also already recalled in the previous cycle (high quality) and those that were only remembered in the final recall (medium quality) in contrast to never-recalled items (low quality). The few items that were remembered in an earlier but not later recall, were excluded from further analyses (see Results and Figure 4). For both age groups, all EEG analyses were conducted on the EEG activity patterns elicited during the first learning phase such that all pairs were novel to the participants and no retrieval-related processes could influence the evoked activity patterns.

### EEG recording and preprocessing

EEG was recorded continuously with BrainVision amplifiers (BrainVision Products GmbH, Gilching, Germany) from 61 Ag/Ag-Cl electrodes embedded in an elastic cap. Three additional electrodes were placed at the outer canthi (horizontal electrooculography (EOG)) and below the left eye (vertical EOG) to monitor eye movements. During recording, all electrodes were referenced to the right mastoid electrode, and the left mastoid electrode was recorded as an additional channel. The EEG was recorded with a pass-band of 0.1 to 250 Hz and digitized with a sampling rate of 1000 Hz. During preparation, electrode impedances were kept below 5 kΩ.

EEG data preprocessing was performed with the Fieldtrip software package (developed at the F. C. Donders Centre for Cognitive Neuroimaging, Nijmegen, The Netherlands; http://fieldtrip.fcdonders.nl; RRID: SCR 004849) and custom MATLAB code (The MathWorks Inc., Natick, MA, USA; RRID: SCR 001622). Data were downsampled to 250 Hz and an independent component analysis was used to correct for eye blink, (eye) movement, and heartbeat artifacts (Jung et al., 2000). Artifact components were automatically detected, visually checked, and removed from the data. For analyses, the EEG was demeaned, re-referenced to mathematically linked mastoids, and band-pass filtered (0.2–100 Hz; fourth order Butterworth). Following the FASTER procedure (Nolan et al., 2010), automatic artifact correction was performed for the remaining artifacts. Excluded channels were interpolated with spherical splines (Perrin et al., 1989). Finally, data epochs of 4 seconds were extracted from −1 s to 3 s with respect to the onset of the scene–word presentation during the study phase (Figure 1A).

**Figure 1:**
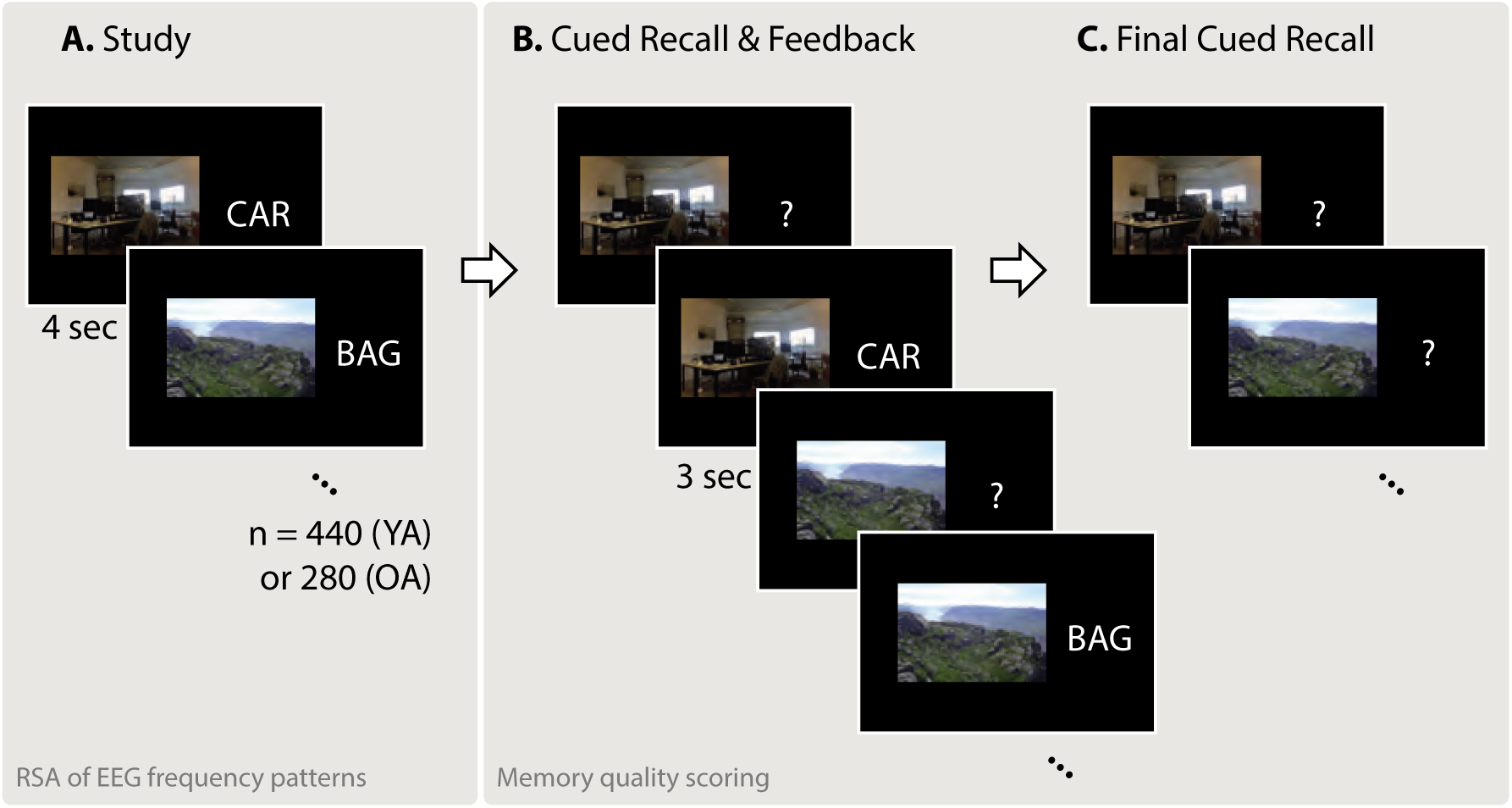
Memory task paradigm (cf. Fandakova et al., 2018). **A.** In the study phase, participants were asked to associate 440 (young adults; YA) or 280 (older adults; OA) scene–word pairs using an imagery strategy. Representational similarity analysis (RSA) was conducted on EEG data during this phase. **B.** During the cued recall and feedback phase, the scene was presented as a cue to verbally recall the associated word. Subsequently, the original pair was presented again for restudy. The cued recall and feedback phase was performed once for younger and twice for older adults. **C.** During final recall, no feedback was provided. Scene–word pairs were sorted into three memory quality categories based on recall performance in phases B and C (see Figure 2).

**Figure 2:**
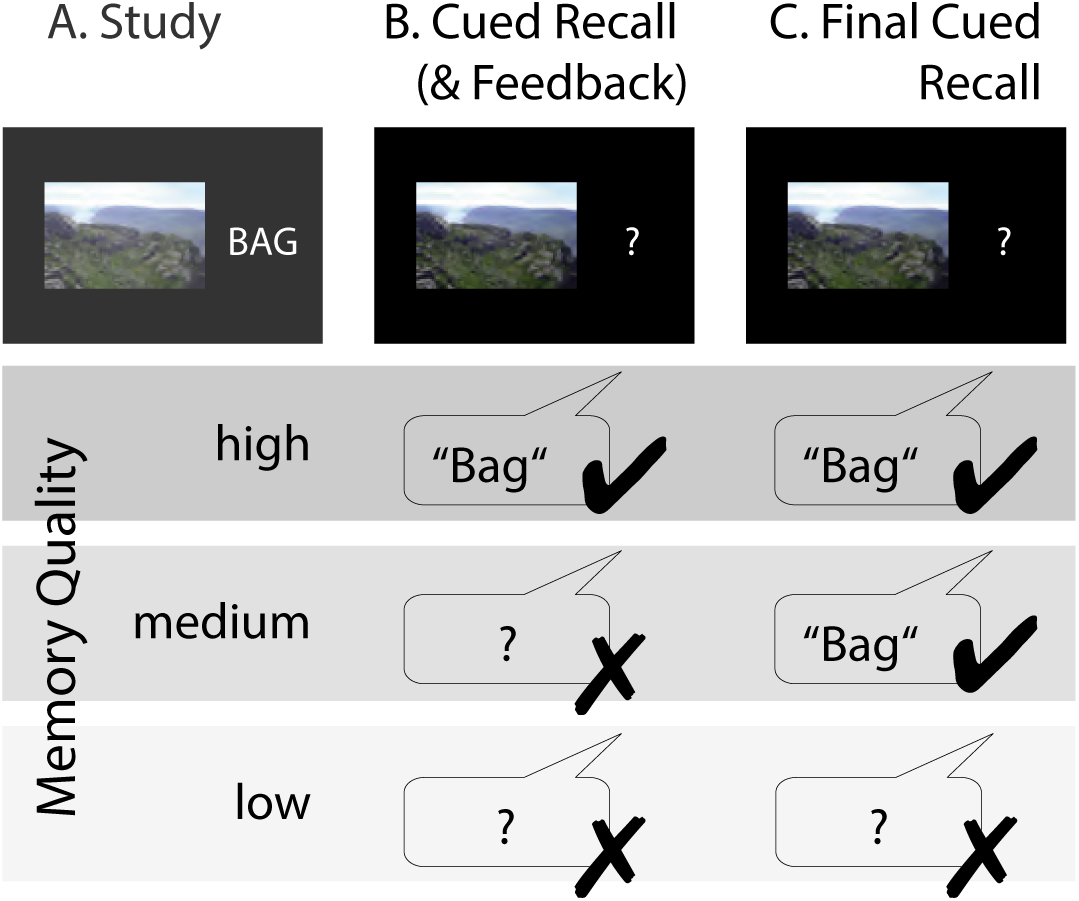
Scoring of stimulus pairs into high, medium, or low memory quality categories based on learning history. For both younger and older adults, items that were correctly recalled in the last recall cycle (C) as well as the previous (B) were scored as high memory quality items. Pairs that were solely recalled in the final recall were scored as medium memory quality items. And items that were never correctly recalled were scored as low memory quality items. Not depicted: Items that were recalled in the earlier but not later recall were excluded. Older adults performed one more cued recall and restudy cycle (between A and B) that was not included in item scoring due to close-to-floor performance. Note that wrong and no responses were treated equally.

### EEG analysis

Time-frequency representations (TFRs) of the data were derived using a multitaper approach. For the low frequencies (2–20 Hz), we used Hanning tapers with a fixed width of 500 ms, resulting in frequency steps of 2 Hz. For higher frequencies (25–100 Hz), we used DPSS (discrete prolate spheroidal sequences) tapers with a width of 400 ms in steps of 5 Hz with seven Slepian tapers resulting in +/-10 Hz smoothing. In this way, we obtained a TFR for each trial and electrode. Trial lengths were reduced to −0.752 s to 3 s relative to stimulus onset.

To counter the effect of intrinsically high correlations between frequency patterns due to the 1/frequency power spectrum (Schönauer et al., 2017), we removed the mean background noise spectrum from the log-transformed TFRs following previously established procedures (i.e., as suggested by the “Better oscillation detection” (BOSC) method; Caplan et al., 2001; Kosciessa et al., 2018; Whitten et al., 2011). Because of structured noise, correlations between different activity patterns are very high and almost never at or below zero, meaning that the true null-distribution is higher than zero. For detailed discussions of these issues (in fMRI), see Allefeld et al. (2016); Cai et al. (2016).

### Multivariate EEG analysis

EEG data were analyzed using representational similarity analysis (RSA; Kriegeskorte et al., 2008). RSA assesses the resemblance of patterns of neural activity, with similar patterns assumed to represent mutual information. In this study, we investigated between-item representational similarity during the first encoding phase in relation to memory quality. “Item” always refers to a scene–word pair. Figure 3 illustrates the procedure for analyzing the similarity between stimulus-specific spatiotemporal frequency representations. RSA was conducted for each participant and EEG channel independently. Stimuli were grouped according to high, medium, and low memory quality (see Figure 2). In order to examine whether between-item representational similarities differed as a function of memory quality, we correlated the noisecorrected and log-transformed frequency patterns of every item with the frequency patterns of all other items of the same memory quality. That is, for each participant we ran three similarity analyses, namely for high, medium, and low memory quality items. In order to use the same number of items for each RSA of a given participant, we reduced them to the number of items available in the condition with the least items. For example, if there were 50 items with high, 180 items with medium, and 210 items with low memory quality for a given participant, the number of items used in the RSAs of medium and low quality items was reduced to 50 as well. Note that the category containing the fewest items was in most cases the group of high memory quality items (except for 6 younger and 6 older participants). We randomly sampled the respective number of items from all available trials of the respective memory quality. As the actual measure of similarity, we employed pairwise Pearson correlations between the corresponding frequency patterns. In each of these correlations, every pair of frequency vectors (with 26 frequency bins) of all time points from the two respective trials were correlated with each other (470 time points, from 752 ms before stimulus onset to 3000 ms after stimulus onset). The resulting time-time similarity matrices were Fisher (z)-transformed. In order to prevent bias towards the randomly picked items, the item sampling was repeated 20 times. Finally, the matrices were averaged to obtain one between-item similarity matrix for each scene–word pair, which indicates the similarity of this pair to all other pairs of the same memory quality. The similarity matrices of all items within one memory quality were then again averaged to obtain the mean similarity matrices between all high, medium, and low memory quality items, respectively. This procedure was performed separately for each of the 60 scalp electrodes.

**Figure 3:**
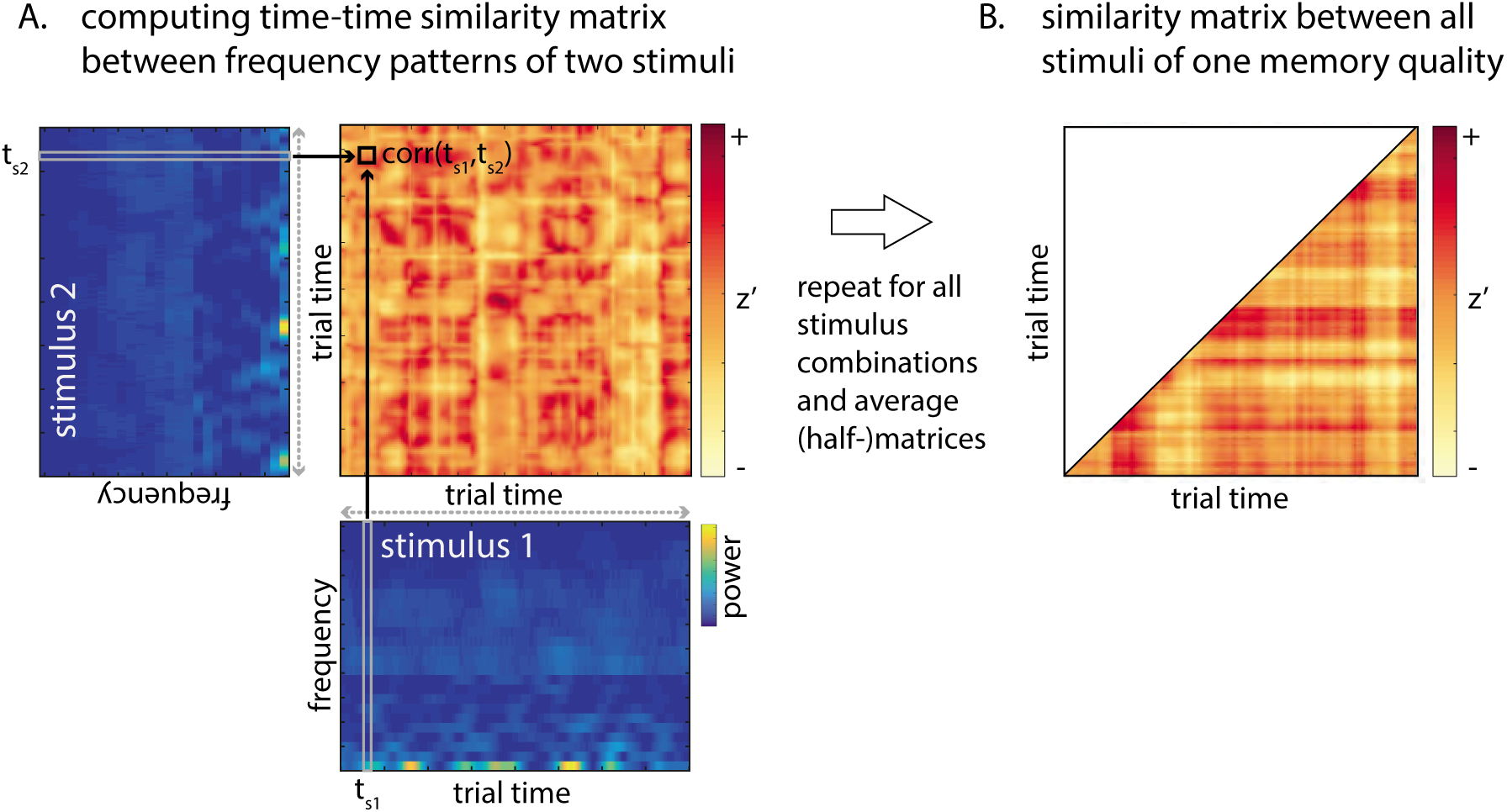
Spectral representational similarity analysis methodology. **A.** The frequency vector from every time point (i.e., column) of the noise-corrected and log-transformed time-frequency pattern (from one electrode) corresponding to stimulus 1 (bottom) is Pearson-correlated with the vectors from every time point of stimulus 2 (left; tilted). For illustration, one example vector of stimulus 1 (t_s1_) and one example vector of stimulus 2 (t_s2_) are highlighted. Correlating these two vectors gives one correlation coefficient, i.e., one coordinate (highlighted with black box) on a matrix with time on both axes. Computing all pairwise time vector correlations results in a time-time similarity matrix representing the similarity of those two frequency patterns at all time point combinations. This procedure is repeated for all items of a certain memory quality (i.e., similarity of stimulus 1 with all others, stimulus 2 with all others, etc.). **B.** Averaging across all similarity matrices yields the mean similarity matrix showing the pattern similarity among all items of the same memory quality. Only one triangle and the diagonal of the matrix are relevant because the similarity of every two frequency patterns is computed twice, resulting in an identical correlation coefficient on both sides of the diagonal. Similarity is quantified as the Fisher *z*-transformed Pearson correlation coefficient (*z*’). Not depicted: This procedure is repeated for all 60 electrodes, the three memory quality categories, and all subjects.

**Figure 4:**
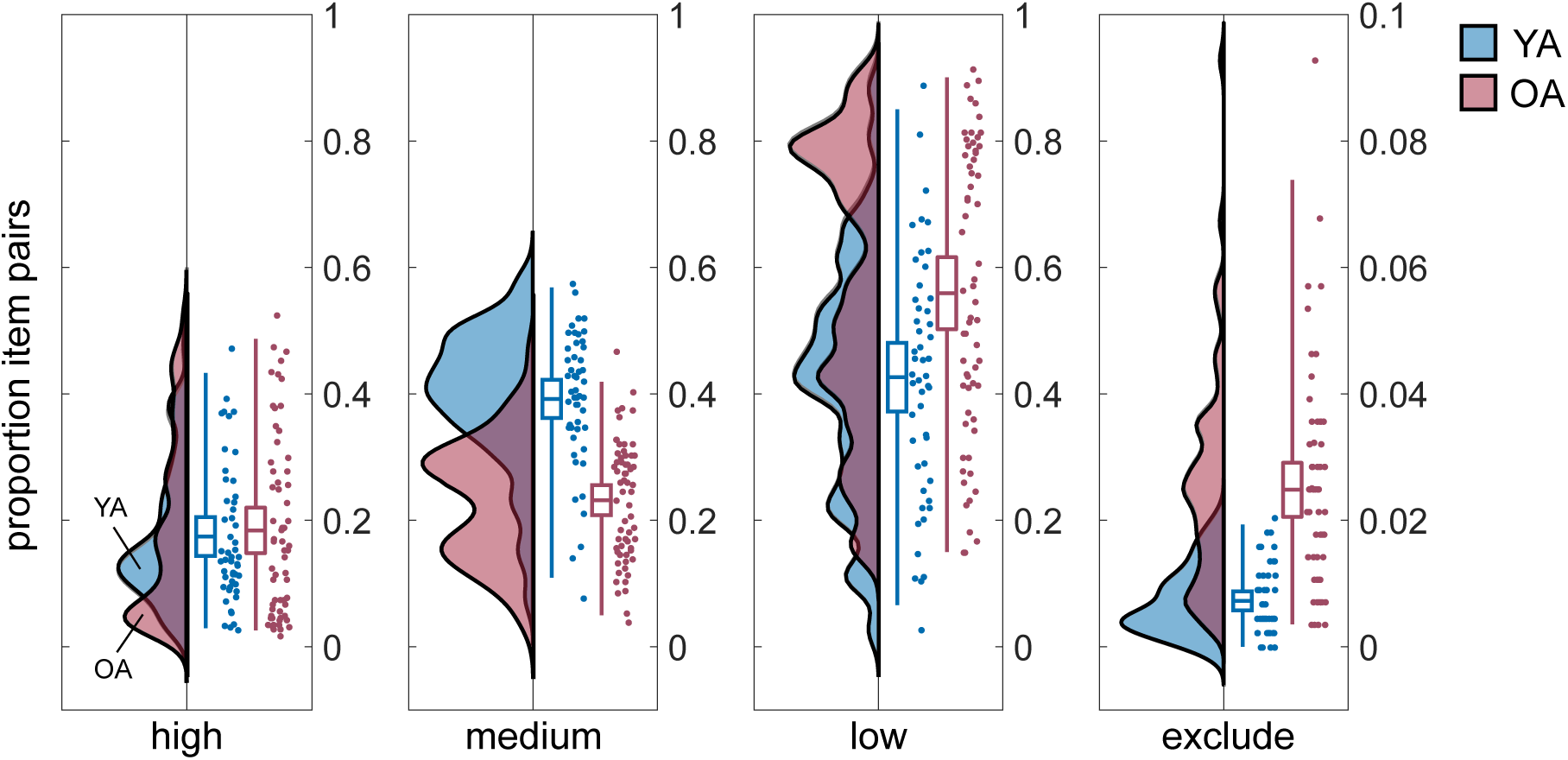
Proportion of item pairs with high, medium, and low memory quality as well as proportion of excluded items for 50 young adults (YA; blue) and 63 older adults (OA; red). Group distributions as un-mirrored violin plots (probability density functions), boxplots with means and 95% confidence intervals, whiskers with 2nd and 98th percentiles, and individual data points (horizontally jittered) (modified from Allen et al., 2018). Note that the y-axis for excluded items differs from that of the other categories. YA studied 440 pairs and OA studied 280 pairs.

The resulting similarity matrices contain the time dimension on both the x- and the y- axis, revealing the frequency pattern resemblance not only at identical within-trial time points (diagonal) but also between all combinations of time points (in analogy to the temporal generalization method; Cichy et al., 2014; King and Dehaene, 2014). This enables us to identify whether certain parts of the memory representations were similar to each other at different times during encoding of the respective scene–word pairs.

Because the similarity of any two items is computed twice and thus the identical correlation coefficients appear twice, namely on both sides of the diagonal, the similarity matrix was reduced to only one of the triangles plus the diagonal.

Representational similarity analyses were computed parallelized on a high-performance computing cluster. All computations and statistics were conducted with Matlab (The MathWorks, Inc., RRID: SCR 001622) versions R2014b or R2016b. The Matlab-based Fieldtrip Toolbox (Maris and Oostenveld, 2007; Oostenveld et al., 2011) (Maris Oostenveld, 2007; Oostenveld et al., 2011; RRID: SCR 004849) was used for performing time-frequency transformations and cluster-based permutation analyses.

### Statistical analysis

#### Memory performance and strategy use

We computed two-sided independent samples *t*-tests in order to test for age differences in the proportion of items within each memory quality category (high, medium, low, forgotten/excluded) and the proportion of items remembered in the final recall task. To compare younger and older adults’ strategy use in the first encoding phase, we used the Wilcoxon rank sum test to examine differences in their median responses of how often they used the imagery strategy.

#### Differences in representational similarity

Within both groups, we tested for differences in the representational similarity matrices between different memory quality categories by conducting non-parametric, cluster-based, random permutation tests (Fieldtrip Toolbox; Maris and Oostenveld, 2007; Oostenveld et al., 2011; RRID: SCR 004849). Univariate two-sided, dependent samples regression coefficient *t*- statistics were calculated for the time-time similarity matrices at all channels. Clusters were formed by grouping neighboring channel *×* time *×* time samples with a *p*-value below 0.05 (spatially and temporally). The respective test statistic was then determined as the sum of all *t*-values within a cluster. The Monte Carlo method was used to compute the reference distribution for the summed cluster-level *t*-values. Samples were repeatedly (100 *×*) assigned into three groups and the differences between these random groups were contrasted to the differences between the three actual conditions (high, medium, and low memory quality). For every repetition the *t*-statistic was computed and the *t*-values summed for each cluster. The *t*-values were *z*-transformed for further analysis.

In addition to the linear regression of all three memory qualities mentioned above, we also compared each pair of memory quality categories using a two-sided, dependent samples *t*-test in the permutation analysis.

We regarded clusters whose test statistic exceeded the 97.5th percentile for its respective reference probability distribution as significant. If such clusters were obtained, we furthermore assessed the time-time intervals and the topographic distributions of the channels showing when and where, respectively, the differences were reliable. The clusters that were identified for each age group were further examined for age and memory quality effects (see below). In addition, we tested for main age group differences in a separate permutation analysis using independent samples *t*-tests.

#### Age and memory quality effects in the identified clusters

To explore potential age differences more closely, we further investigated the relationship between pattern similarity and memory quality by conducting independent samples regression coefficient *t*-statistics for each participant. We then extracted and averaged the individual ( z)transformed regression coefficients within the time-time-electrode clusters that were identified in younger and older adults (see above). For both clusters and age groups we performed one-sample *t*-tests to test whether the correlation coefficients come from a distribution with a mean different from zero. Furthermore, we tested for differences between the age groups in both clusters using independent samples *t*-tests.

## Results

### Memory performance and strategy use

During the cued recall phases, participants had to respond verbally with the word they previously learned to associate with the presented image. We sorted the trials according to whether recall was successful, and when, into high, medium, and low memory quality items (see Methods). The proportion of high memory quality items did not differ between younger adults and older adults (*M* (younger adults) = 0.17, SD(younger adults) = 0.11, *M* (older adults) = 0.18, SD(older adults) = 0.15; *t* (111) = −0.4, *p* = 0.69, two-sample *t*-test; see Figure 4). In contrast, the proportion of items with medium memory quality was significantly larger for younger than older participants (*M* (younger adults) = 0.39, SD(younger adults) = 0.11, *M* (older adults) = 0.23, SD(older adults) = 0.09; *t* (111) = 8.48, *p* = 10^-13^), while older adults had a significantly higher proportion of low memory items (*M* (younger adults) = 0.43, SD(younger adults) = 0.19, *M* (older adults) = 0.56, SD(older adults) = 0.23; *t* (111) = −3.31, *p* = 0.0012). Note that in older adults we observed a higher proportion of items that were remembered in an early but not later recall phase, i.e., that were forgotten in the course of the experiment (*M* (younger adults) = 0.007, SD(younger adults) = 0.005, *M* (older adults) = 0.025, SD(older adults) = 0.02; *t* (111) = −7.04, *p* = 1.6 *×* 10^-10^). Those item pairs were excluded from further analyses.

Our experimental procedure was successful in inducing variability in memory performance such that both groups could remember approximately half of the studied items: Young adults successfully recalled on average 56.64 % (SD = 10.7) and older adults successfully recalled on average 41.6 % (SD = 12.06) of the items (440 and 280, respectively). However, our procedure did not completely eliminate age differences since young adults still performed significantly better than older participants in the final recall task (*t* (111) = 3.82, *p* = 0.0002, two-sample *t*-test).

After the first study phase, we asked participants to indicate on a four-point scale how often they had used specific memory strategies for the task (1: almost always, 4: almost never). With regard to the imagery strategy, young adults indicated that they used it significantly more often than older adults (younger adults: median = 1.5, min = 1, max = 3; older adults: median = 2, min = 1, max = 4; *z* = −5.09, *p* = 0.0000004, Wilcoxon rank sum test).

### Representational similarity

Calculation of between-item representational similarity was based on the initial encoding phase (Figure 1A). To identify whether high pattern resemblance or high pattern distinctiveness during learning was beneficial for later memory success, we sorted all items according to subsequent memory performance and correlated the evoked spatiotemporal frequency pattern of each item with every other item in the same memory quality category. The resulting mean similarity matrices over all channels and scene–word pairs are shown in Figure 5A. These matrices display the similarity of the frequency representations at all possible within-trial time point combinations (−0.752 s to 3 s relative to stimulus onset at 0). In contrast, the diagonals of the similarity matrices (also plotted separately in Figure 5B) only show the similarity between items at identical time points and facilitate a visual comparison of the time courses of representational similarities for the different memory quality categories and age groups. Although this omits much of the similarity information, elevated similarities do occur largely along the diagonal. Note that the diagonals are only plotted for illustration purposes and all statistical tests were performed on the complete matrices as presented in Figure 5A.

**Figure 5:**
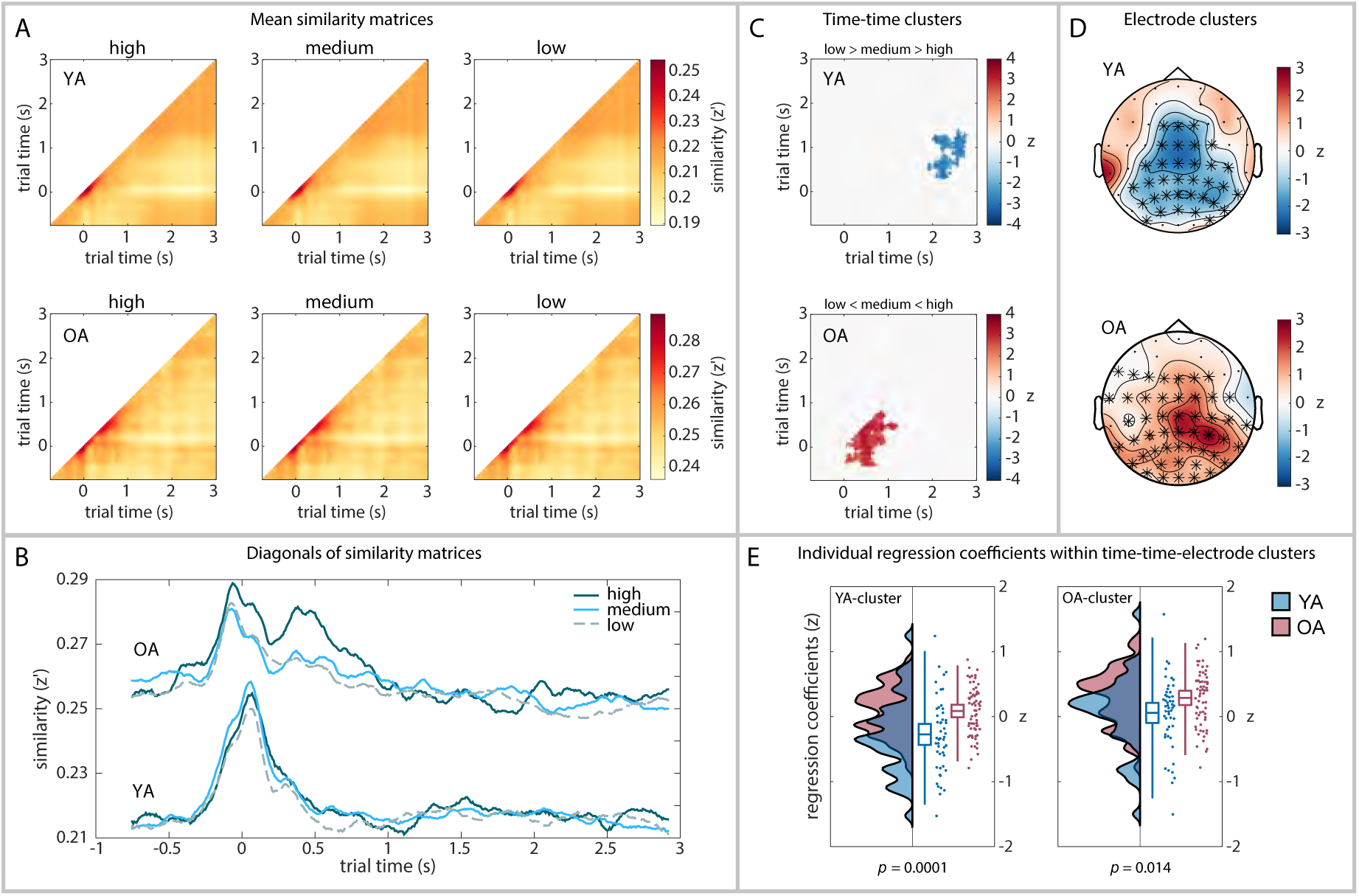
Between-item pattern similarities and statistics. Similarity is quantified as Fisher *z*-transformed Pearson correlation coefficient (*z*’). On time axes, zero denotes stimulus onset. C and D show results from cluster-based permutation analyses for each age group, E shows results from individual regression analyses (see Methods). **A.** Mean time-time similarity matrices across all 60 channels and items within each memory quality category (high, medium, low) for all 50 young adults (YA; top) and 63 older adults (OA; bottom). Note that the scales differ between age groups. **B.** Diagonals from the time-time similarity matrices (see A). **C.** Time-time clusters (masked *z*-scores) in which the three memory quality categories differ significantly within each age group (averaged across reliable electrodes, see D). **D.** Topographic representations of the electrode clusters that revealed reliable differences between memory quality categories within each age group (averaged across reliable time windows, see C). **E.** *Z*-transformed regression coefficients extracted from time-time-electrode clusters identified in YA (left) and OA (right) (see C and D). Group distributions (probability density functions), boxplots with means and 95% confidence intervals, whiskers with 2nd and 98th percentiles, and individual data points (horizontally jittered) for YA (blue) and OA (red) (modified from Allen et al., 2018). ( P)-values are given for group differences within each cluster (independent samples *t*-tests). *Note the difference between z’ (Fisher z-transformed correlation coefficients) and z (z-transformed regression coefficients).*

#### Older adults exhibit generally higher representational similarity than young adults

Shortly before stimulus onset, similarity increased in both age groups and reached a peak around the time of onset (Figure 5A,B). Elevated similarity occurred mainly between identical trial time points (diagonal) with slightly more persistent activity (elevated off-diagonal similarity) in older adults compared to young adults. Irrespective of later memory success, between-item pattern similarity was generally higher in older adults than in young adults (averaged across the whole time-time matrix and all 60 channels: *M* (younger adults) = 0.21, SD(younger adults) = 0.065, *M* (older adults) = 0.25, SD(older adults) = 0.068; 500 cluster permutations, *p* = 0.002).

#### Representational similarity differentially affects memory performance of younger and older adults

Within both age groups, we tested for differences in the levels of representational similarity between scene–word pairs of different memory quality by conducting linear regressions. We controlled for multiple comparisons by using non-parametric cluster-based permutation tests. In both age groups we identified a cluster with a Monte Carlo *p*-value below 0.025, which indicates a reliable linear relationship between representational similarity and memory quality (young adults: *p*=0.0099; older adults: *p*=0.0099; see Figure 5C). Importantly, the direction of this relationship differed between groups: while the relation between similarity and memory quality was positive in older adults (low *<* medium *<* high), it was negative in young adults (low *>* medium *>* high) (Figure 5E).

The cluster obtained in older adults included most of the diagonal from 50 ms to 830 ms after stimulus onset and extended off-diagonally to 470 ms before and 1240 ms after stimulus onset (Figure 5C). Elevated similarity along the diagonal indicates similarity between neural representational patterns at identical trial time points, whereas off-diagonal time windows suggest similar activation patterns at different trial time points. The larger the distance of a coordinate from the diagonal, the more distant are the compared time points in the respective frequency patterns. Differences between memory quality categories were reliable in most (49 out of 60) occipital, parietal, temporal, and central electrodes in older adults (Figure 5D).

In contrast to the cluster found in older adults, an off-diagonal cluster was identified for young adults, in which low memory quality items displayed significantly more similarity than medium and high memory quality items (Figure 5C). Compared to older adults, where differences between memory qualities were found to be most pronounced between early and neighboring trial time points, i.e., close to the diagonal, the off-diagonal cluster identified in young adults indicated that differences occurred at later and more distant trial time points. Specifically, differences were found between earlier (450 ms to 1400 ms after stimulus onset) and later time points (2640 ms to 2800 ms after onset) and at 34 mainly parietal-occipital and central electrodes (Figure 5D). Despite the relatively poor spatial resolution in EEG, the large electrode clusters in both young and older adults indicate that memory representations are broadly distributed over the brain rather than specific to a particular region.

Additional analyses of pairwise comparison of the three memory quality categories instead of linear regression resulted in the same significant cluster only for high versus low quality items in older adults, and no significant differences among memory quality categories in young adults.

#### Age and memory quality effects in the identified clusters

The cluster-based analyses reported above suggested differential memory-related representational similarity in younger and older adults. To explore potential age differences more closely, we additionally tested for a linear relationship between representational similarity and memory quality in each participant by conducting individual linear regressions. We then extracted and averaged the individual *z*-transformed regression coefficients within each timetime-electrode cluster (see Figure 5E). In the young-adult cluster only the mean regression coefficients of the young adults differed from zero (young adults: *t* (49) = −3.42, *p* = 0.0013; older adults: *t* (62) = 1.79, *p* = 0.08; one-sample *t*-tests) and vice versa in the older-adult cluster (young adults: *t* (49) = 0.75, *p* = 0.46; older adults: *t* (62) = 5.27, *p* = 0.000002). In both clusters the regression coefficients differed significantly between younger and older adults (young-adult cluster: *M* (young adults) = −0.27, SD(young adults) = 0.57, *M* (older adults) = 0.086, SD(older adults) = 0.38, *t* (111) = −4.03, *p* = 0.0001; older-adults cluster: *M* (young adults) = 0.058, SD(young adults) = 0.55, *M* (older adults) = 0.29, SD(older adults) = 0.43, *t* (111) = −2.5, *p* = 0.014; independent samples *t*-tests) implying that age differences do exist in the relation between representational similarity and memory quality in these clusters.

## Discussion

The present study aimed to reconcile an evident tension between theories relating neural pattern similarity and memory in the fields of cognitive neuroscience and cognitive aging research. We addressed the central question whether high pattern resemblance or high pattern distinctiveness benefits memory performance. To this end, we computed the similarity between the EEG frequency patterns elicited during encoding of different scene–word pairs at each electrode and related this measure of between-pair similarity to subsequent recall performance of younger and older adults.

For older adults, between-item representational similarity was generally higher compared to young adults, supporting the “dedifferentiation” hypothesis of declining neural distinctiveness with age (Baltes and Lindenberger, 1997; Carp et al., 2011; Li et al., 2004; Park et al., 2004; Park et al., 2012; Payer et al., 2006; St-Laurent et al., 2014). Previous studies suggested that the loss of neural specificity in old age may underlie age-related cognitive impairments. This was, for example, supported by the finding that the level of neural distinctiveness and fluid intelligence were correlated (Park et al., 2010). However, most previous studies were not able to directly link neural item specificity with study participants’ performance since memory for the items themselves was not assessed. By measuring between-item representational similarity during the encoding phase of an associative memory task and sorting the trials according to subsequent memory performance, we were able to directly relate measures of neural distinctiveness during encoding to later recall success.

Specifically, based on learning history, we sorted the studied scene-word pairs into high, medium, and low memory quality items and, on the within-subject level, measured the linear relationship between the level of representational similarity and memory quality. Importantly, the direction of this relationship as well as the time window in which representational similarity mattered for subsequent memory performance differed between younger and older participants: For older adults, *high* similarity early during encoding (470 ms before stimulus onset to 1240 ms after stimulus onset) benefited memory performance. For young adults, *low* similarity between earlier (450 ms to 1400 ms after stimulus onset) and later time points during encoding (2640 ms to 2800 ms after onset) benefited memory performance.

That is, although older adults remembered significantly fewer items and revealed overall higher between-item similarity than younger adults, on the within-subject level, item representations with high similarity to other items were actually those that older adults remembered best. Hence, while the age group differences replicated previous reports of increased neural similarity in older compared to younger adults, the within-person direction of the similaritymemory association among older adults corroborates cognitive models of memory (Clark and Gronlund, 1996; Gillund and Shiffrin, 1984; Hintzman, 1988) as well as previous memory studies with younger adults. These studies showed that higher similarity between different item representations (often called ‘global similarity’) is beneficial for subsequent recognition memory (LaRocque et al., 2013; Lu et al., 2015; Ye et al., 2016), memory confidence and categorization (Davis et al., 2014), fear memory (Visser et al., 2013), and associative memory formation (Wagner et al., 2016).

FMRI experiments located this beneficial effect of representational similarity in medial temporal lobe regions, whereas in the hippocampus, pattern distinctiveness supported memory (LaRocque et al., 2013). Indeed, impaired pattern separation computations in the hippocampus were reported for older adults (Shing et al., 2011; Wilson et al., 2006; Yassa et al., 2011). While high pattern distinctiveness may be beneficial for memory performance to prevent false memories, high pattern similarity may support mnemonic decisions by capturing regularities across experiences (LaRocque et al., 2013) and by giving rise to a feeling of familiarity (Davis et al., 2014). Higher pattern similarity may also reflect more consistent processing that facilitates associative memory formation (Wagner et al., 2016). Strikingly, a tendency for more generalized memories (Koutstaal and Schacter, 1997; Koutstaal et al., 2001; Tun et al., 1998) and a stronger reliance on familiarity (Light et al., 2000; Prull et al., 2006; Yonelinas, 2002) is indeed often reported for older adults. Our findings suggest that these behavioral patterns result from an overall increased neural similarity.

Surprisingly, although for older adults items that were successfully learned also showed higher pattern similarity, we did not identify this beneficial effect of pattern similarity in young adults. Given that most of the studies that reported this effect in young adult samples tested recognition memory (Davis et al., 2014; LaRocque et al., 2013; Lu et al., 2015; Ye et al., 2016), the benefit may be less pronounced in (cued) recall tasks (but compare (Wagner et al., 2016) who used a picture–location association task). Whereas a sense of familiarity as a consequence of high pattern similarity (Davis et al., 2014; Gillund and Shiffrin, 1984) can be sufficient for successful recognition, recall typically requires retrieval of specific details of the studied items (Craik and Tulving, 1975). Therefore, the beneficial effects of high pattern similarity may be identified more easily in pure recognition memory tasks and/or participant groups who base their mnemonic decisions more strongly on familiarity signals, such as older adults.

The observed age group differences are in line with previous suggestions that external stimuli exert a stronger drive on neural processing in older than in younger adults (Lindenberger and Mayr, 2014; Sander et al., 2012; Werkle-Bergner et al., 2012). In line with the “load-shift” model of executive functioning in aging (Velanova et al., 2007), the high, externally triggered similarity of scene–word pairs may have helped older adults to memorize pairs based on familiarity. At the same time it may have impaired their ability to form differentiated mnemonic representations early on. The resulting burden on late selection processes might have impaired older adults’ ability to engage elaboration mechanisms supporting the formation of distinctive memories. By contrast, the advantage of reduced similarity in younger adults may hint at their ability to engage elaborative mechanisms supporting future recall of detailed mnemonic information, as observed in the off-diagonal effect. In sum, we suggest that older adults’ advantage of high between-pair representational similarity early in the trial may stem from a reliance on familiarity-based remembering, while younger adults exploited more recall-based strategies, capitalizing on a higher capacity to form discrete representations later in the trial. We would like to speculate that the benefit of distinct neural activation patterns is especially prominent in the deployed task, in which participants were explicitly instructed to form very distinct mental images of the corresponding scene–word pair. Although older adults were extensively trained in using the imagery technique of forming salient mental images that integrate the associated picture and word, the post-encoding strategy questionnaire showed that they utilized this strategy less frequently than young adults. This may explain their lower recall performance despite having studied fewer pairs and having more opportunity to rehearse them. This conjecture is supported by previous research showing that older adults continue to use other mnemonic strategies even though they have learned about the benefits of imagery (Hertzog et al., 2012).

So far, the prevailing available evidence on the relationship between representational similarity and memory performance has been based on fMRI studies and therefore lacks insights into the temporal dynamics of pattern similarity during the formation of memory representations. Here, we demonstrate the advantage of dissociating different parts within the trial time course that reveal distinctions in the way representational similarity relates to memory performance of younger and older adults.

An open question is how between-item similarity links to item-specific representational stability (across item repetitions or between encoding and retrieval). Recent research suggests that representational stability benefits memory performance (Lu et al., 2015; Xue, 2018; Xue et al., 2010) and declines in old age (St-Laurent et al., 2014; Zheng et al., 2017). Understanding the mutual influences of between-item similarity and representational stability may be crucial to complete our comprehension of how memories are represented in the brain across the lifespan.

In summary, we provide critical new evidence that the often observed between-subject effect of generally higher similarity between neural representations in older adults does not predict their future memory success besides the fact that they perform worse than young adults who exhibit generally lower pattern similarity. Instead, on the within-subject level, older adults best remembered the items with the highest peak in pattern similarity early du ring encoding. Moreover, we show that young adults benefited from eliciting distinct memory representations later during the encoding trial, which presumably reflects the implementation of the imagery strategy for scene–word binding. The work presented here extends our knowledge about between-item pattern similarity as a memory-relevant representational property. In particular it shows how its relation to cognitive performance may change in the course of aging.

## Acknowledgements

This study was conducted within the “Cognitive and Neural Dynamics of Memory across the Lifespan” (CONMEM) project at the Center for Lifespan Psychology, Max Planck Institute for Human Development. The research was partially financed by the Max Planck Society. MWB’s work was supported by a grant from the German Research Foundation (DFG, WE 4269/3-1, YLS as Co-PI) as well as an Early Career Research Fellowship 2017–2019 awarded by the Jacobs Foundation. YLS and MCS were each supported via Minerva Research Groups awarded by the Max Planck Society. YLS is funded by the European Union (ERC-2018-StG-PIVOTAL-758898) and a Fellowship from the Jacobs Foundation (JRF 2018-2020). VRS is a fellow of the International Max Planck Research School on the Life Course. We thank Beate Mühlroth and Xenia Grande for organizing data collection, Kristina Günther for help in participant recruitment, Julia Delius for editorial assistance, Michael Krause for help with cluster computing, and all student assistants who helped with data collection. We are grateful to all members of the CONMEM project for helpful feedback on the analysis. Finally, we thank all study participants for their time.

